# Refractive index tomograms and dynamic membrane fluctuations of red blood cells from patients with diabetes mellitus

**DOI:** 10.1101/087460

**Authors:** SangYun Lee, HyunJoo Park, Kyoohyun Kim, YongHak Sohn, Seongsoo Jang, YongKeun Park

## Abstract

In this paper we present the optical characterisations of diabetic red blood cells (RBCs) in a non-invasive manner employing three-dimensional (3-D) quantitative phase imaging. By measuring 3-D refractive index tomograms and 2-D time-series phase images, the morphological (volume, surface area and sphericity), biochemical (haemoglobin concentration and content) and mechanical (membrane fluctuation) parameters were quantitatively retrieved at the individual cell level. With simultaneous measurements of individual cell properties, systematic correlative analyses on retrieved RBC parameters were also performed. Our measurements show that diabetic patients had RBCs of reduced cell sphericity and elevated intracellular haemoglobin concentration and content compared to healthy (non-diabetic) subjects. Furthermore, membrane deformability of diabetic RBCs is significantly lower than that of healthy, non-diabetic RBCs. Interestingly, non-diabetic RBCs exhibit strong correlations between the elevated glycated haemoglobin in RBC cytoplasm and decreased cell deformability, whereas diabetic RBCs do not show correlations. Our observations strongly support the idea that slow and irreversible glycation of haemoglobin and membrane proteins of RBCs by hyperglycaemia significantly compromises RBC deformability in diabetic patients.

## Introduction

Diabetes mellitus is a life-threatening disease affecting many people. Consequences of all diabetes types include chronically elevated blood glucose levels, i.e. hyperglycaemia, caused by a lack of insulin or loss of insulin-mediated metabolic functions, which eventually results in a wide range of complications ranging from microvascular diseases including retinopathy, neuropathy and nephropathy, and cardiovascular diseases, to severe depression and dementia^1^.

Characterisation of the deformability of red blood cells (RBCs) is crucial to understanding the pathophysiology of diabetes because it is strongly related to the disease’s complications. The deformability of diabetic RBCs has been extensively investigated using various experimental approaches including optical microscopy^2^, Coulter counter^3^ and laser diffraction ektacytometry^4^. In particular, the main cytoplasmic alteration in diabetic RBCs caused by hyperglycaemia, the glycation of haemoglobin (Hb), has gained increasing attention because the relative amount of glycated Hb, mainly consisting of HbA1c to total Hb, reflects the mean plasma glucose concentration of the previous three months^5^. Also, the glycosylation of other intracellular and membrane proteins of diabetic RBCs has been reported^6^. Furthermore, damage to RBC membranes resulting from chronic hyperglycaemia have been accessed by measuring end products of lipid-peroxidation^7, 8^. It has been known that diabetic RBCs exhibit significantly affected deformability compared to non-diabetic RBCs using micropipette aspiration^9^, ektacytometry^10^, fabrication of micro-channels^11^ and membrane filters mimicking capillary vessels^12^. Instant effects of *in vitro* glucose treatments on RBC deformability have been addressed^13^.

These investigative approaches have significantly enhanced our understanding of the rheology of diabetic RBCs. However, none of these approaches can simultaneously probe morphological, biochemical and mechanical parameters, particularly at the individual cell level. Moreover, such previous approaches depend on large external loads or deformation of RBCs via physical contact, and thus are not well suited to precise measurement of soft and elastic properties of RBCs within linear deformation regimes. Accordingly, previous approaches do not allow systematic correlative analyses to be performed on simultaneously measured various cellular parameters. As a consequence, a systematic study on the integrated effects of diabetic complications or hyperglycaemia on individual human RBCs has not been fully investigated previously.

To effectively address the existing problems, we present a systematic study measuring individual parameters in RBCs from patients with diabetes using the quantitative phase imaging (QPI) technique^14, 15^. QPI is an interferometric imaging technique which can precisely and quantitatively measure live cells and tissues, utilising the refractive index (RI) as an intrinsic optical contrast. Recently, QPI techniques have been utilised for the study of various biological and medical areas, including malarial infection^16-18^, haematology^19, 20^, neuroscience^21^ and sickle cell disease^22, 23^. Using the QPI technique, we performed label-free and quantitative measurements of live RBCs, and assessed three-dimensional (3-D) refractive index (RI) tomograms and 2-D time-series phase maps of individual cells. From the measured 3-D RI tomograms and the time-series phase images, the morphological (volume, surface area and sphericity), biochemical (Hb concentration and Hb content) and mechanical (membrane fluctuation) parameters were retrieved. Also, correlative analyses among six measured RBC parameters were systematically performed and analysed with the information about independently measured HbA1c.

## Results

To reconstruct the 3-D morphologies of individually measured RBCs, we employed a QPI technique called common-path diffraction optical tomography (cDOT)^24, 25^ (see Materials and Methods). Employing the principle of optical diffraction tomography^26-28^, cDOT measures 3-D RI tomograms of cells and tissues, which is optically analogous to X-ray computed tomography. Multiple 2-D holograms of a sample were recorded with various angles of laser illumination, from which complex optical fields of light at the sample plane were retrieved using a field retrieval algorithm^29^. Then, from a set of the retrieved multiple 2-D optical fields, a 3-D RI distribution of the sample was reconstructed.

Using cDOT, we measured 3-D RI tomograms of 40 RBCs per person for a total of 12 non-diabetic and diabetic blood donors (see Materials and Methods). The reconstructed 3-D RI tomograms and the RI isosurfaces (〈*n*〉 > 1.35) of the representative non-diabetic and diabetic RBCs are presented in Figure 1. Cross-sectional images of the measured RI tomograms of both the non-diabetic and diabetic RBCs show characteristic biconcave shapes. Because the RI value of the RBC cytoplasm is linearly proportional to the intracellular Hb concentration with a known proportionality coefficient, RI increment^30^, the RI value of RBC can be directly related to Hb concentration.

**Fig. 1.**
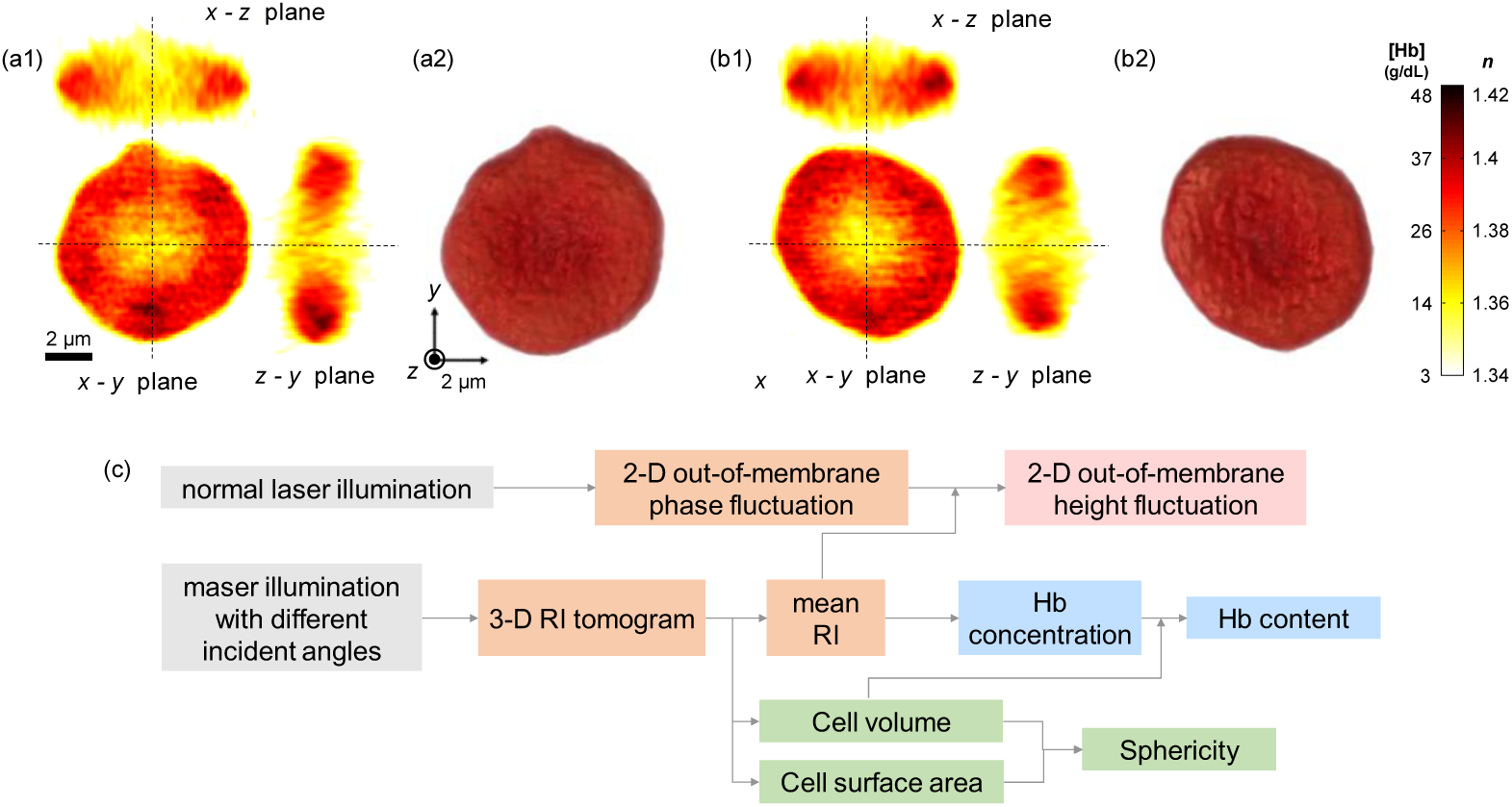
(a–b) Reconstructed 3-D RI tomograms and isosurface renderings of representative non-diabetic and diabetic RBCs. Cross-sectional images (along the *x*-*y*, the *z*-*y* and the *x*-*z* planes) of the 3-D RI tomogram of the representative non-diabetic (a1) and diabetic (b1) RBCs. RI isosurfaces (*n* > 1.35) of the corresponding non-diabetic (a2) and diabetic (b2) RBCs were presented. (c) Flow diagram for retrieval of six RBC parameters.

Figure 1(c) summarises the procedure required for retrieving quantitative RBC parameters (see Materials and Methods). The morphological parameters (cell volume and surface area) are directly obtained from the reconstructed 3-D RI tomogram. Sphericity is a dimensionless number, defined as a normalised volume-to-surface area ratio, which addresses the sphere-likeness of cells. Perfect spheres and flat disks have a sphericity of 1 and 0, respectively. Cytoplasmic Hb concentration of RBCs is extracted from the measured RI value because RBC cytoplasm mainly consists of Hb solution, and there is a linear relationship between the RI of the Hb solution and its Hb concentration^30, 31^. Then, the retrieved Hb concentration and cell volume offer information about the cytoplasmic Hb content. While laser illumination is set to be perpendicular to an RBC, optical phase delays contributed from the cell membrane fluctuation can be continuously recorded using a high-speed camera. Finally, dynamic membrane fluctuations of individual cells are calculated from 2-D phase fluctuations and mean RI values from the reconstructed 3-D RI tomograms. Recently, it has been reported that these RBC fluctuations are acutely affected by diverse pathophysiological conditions^17,18,22,32-35^.

### Retrieved RBC parameters of all non-diabetic and diabetic blood donors

To address the morphological and biochemical alterations in human RBCs by diabetes mellitus, we retrieved three morphological (volume, surface area and sphericity) and two biochemical (cytoplasmic Hb concentration and Hb content) RBC parameters for every measured RBC from its 3-D RI tomogram.

As shown in Figure 2(a), there is no statistical difference in volume distributions of non-diabetic and diabetic RBCs. The mean volumes of non-diabetic and diabetic RBCs are 90.5 ± 11.4 [mean ± standard deviation (SD)] and 90.2 ± 10.9 fL, respectively. Diabetic RBCs exhibit slightly higher surface area compared to that of non-diabetic RBCs (*p*-value < 0.01). The mean surface areas of non-diabetic and diabetic RBCs are 144.1 ± 17.4 and 148.2 ± 14.4 μm^2^, respectively. The calculated mean sphericity values of non-diabetic and diabetic RBCs are 0.68 ± 0.06 and 0.66 ± 0.05, respectively. The enlarged membrane surface area of diabetic RBCs mainly accounts for these significant diminutions of RBC sphericity.

**Fig. 2.**
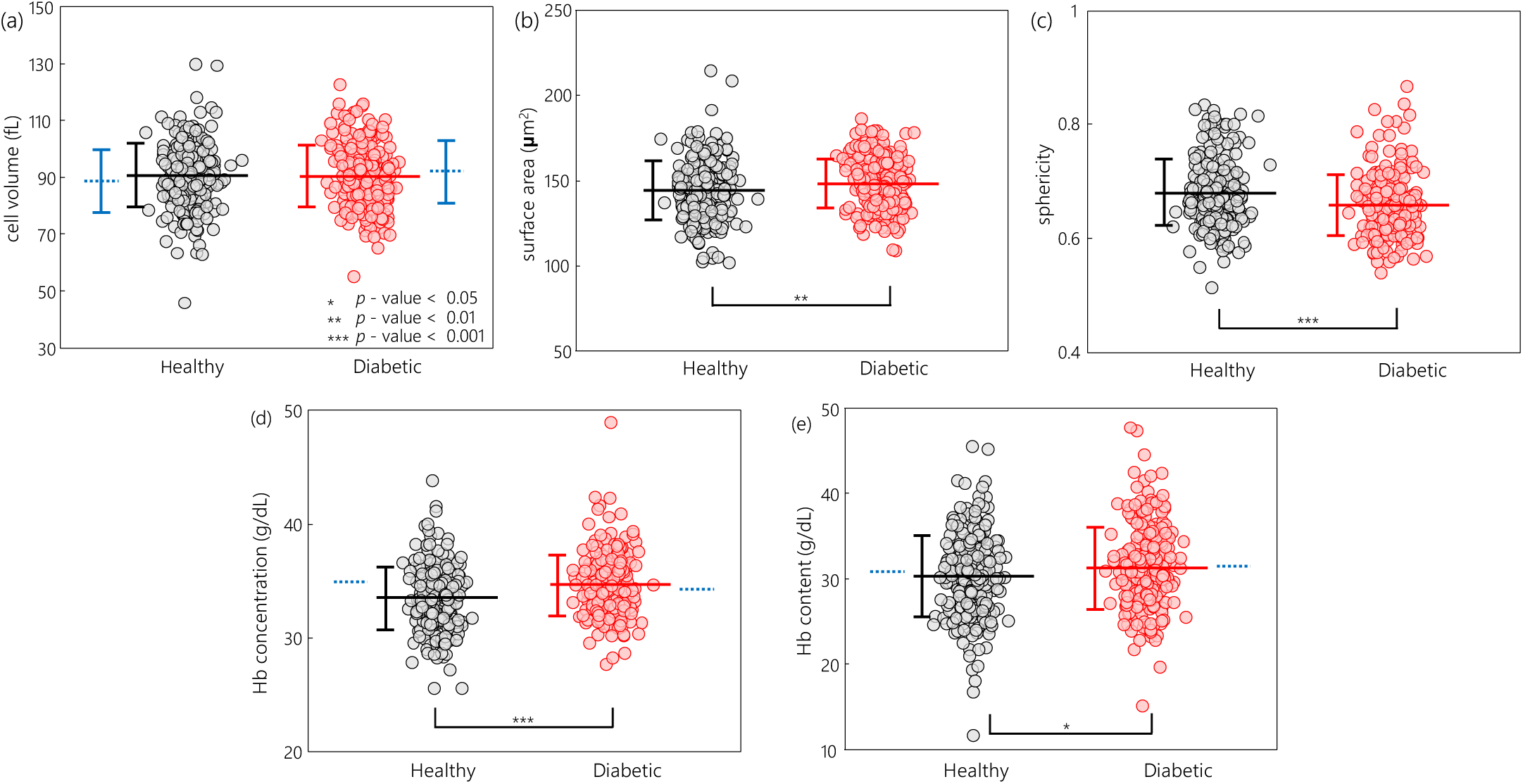
Morphological and biochemical parameters for all measured RBCs from healthy controls and diabetic patients: (a) volume, (b) surface area, (c) sphericity, (d) cytoplasmic Hb concentration and (e) Hb content. Each circle denotes an individual RBC measurement. The horizontal lines and the error bars represent the calculated mean value and sample standard deviation, respectively. The blue dotted lines correspond to results from complete blood count tests.

The retrieved Hb concentration and the Hb content of non-diabetic and diabetic RBCs are presented in Figures 2(d) and (e), respectively. The diabetic RBCs exhibit significantly elevated cytoplasmic Hb concentration as compared to the non-diabetic RBCs (*p*-value < 0.001). The calculated mean Hb concentration of non-diabetic and diabetic RBCs are 33.4 ± 2.8 and 34.6 ± 2.7 g/dL, respectively. The cytoplasmic Hb content of diabetic RBCs is also slightly higher than the control RBCs, with a *p*-value of about 0.03. The retrieved mean Hb content in healthy and diabetic RBC cytoplasm is 30.3 ± 4.8 and 31.2 ± 4.8 pg, respectively. Our optical measurements on cytoplasmic Hb content agree well with the mean corpuscular Hb values of blood donors which were independently measured using the complete blood count machine (the blue dotted lines in Figure 2). The main cause of observed light increase in Hb content of diabetic RBCs is thought to be caused by the additional formation of stable ketoamine linkages on Hb molecules by glucose through a non-enzymatic, slow glycation process (ref). The retrieved RBC parameters of individual subjects with both complete blood count (CBC) result sheet and independently determined HbA1c level are presented in the *Supplementary Information* (see *Supplementary Figure S1*).

### Alterations in mechanical property of RBCs by diabetes mellitus

Cell deformability represents the ability of a cell to change and restore its shape in response to external stimuli. In particular, the remarkably high deformability of RBCs has a core role in transporting oxygen to body organs and tissues through capillaries with diameters even smaller than those of RBCs. It has been widely accepted that the main factors governing the overall RBC deformability include the cell sphericity, mechanical properties of membranes and cytoplasmic viscosity^36, 37^, which are mainly affected by alterations in cell volume or surface area, cytoskeletal spectrin networks and intracellular Hb concentrations, respectively.

To investigate the mechanical alterations in RBC membranes by diabetes, we measured dynamic membrane fluctuations of non-diabetic and diabetic RBCs. Recently, it has been shown that QPI techniques can be used to precisely probe dynamic fluctuations in cell membranes^38^, and these have been utilised in diverse applications in infectious disease^17, 35^, sickle cell disease^39^ and haematology^19,40,41^.

The 2-D topographic height maps of representative non-diabetic and diabetic RBCs are shown in Figures 3(a) and (b), respectively. The corresponding dynamic membrane fluctuation maps, defined as temporal SDs of the RBC height profiles, are also presented in Figures 3(c) and (d) (Materials and Methods).

**Fig. 3.**
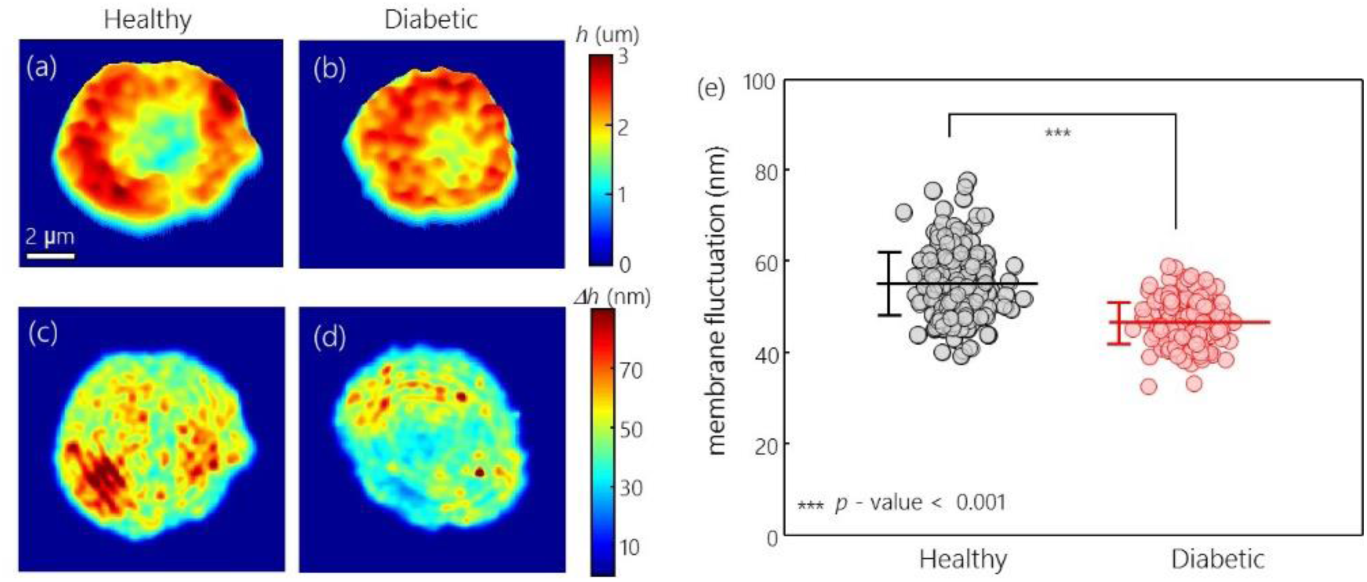
2-D membrane height maps of the representative RBCs from (a) healthy controls and from (b) diabetic patients, respectively. 2-D membrane fluctuation maps of the corresponding RBCs are represented in (c) and (d). (e) Graph for retrieved membrane fluctuations of every measured RBC. The horizontal lines and the vertical error bars denote the mean value and sample standard deviation of measured membrane fluctuations, respectively.

To investigate cell deformability, we spatially averaged the 2-D membrane fluctuation map of individual RBCs and defined it as the mean membrane fluctuation. As shown in Figure 3(e), it is clear that diabetic RBCs exhibit significantly decreased mean membrane fluctuations compared to healthy RBCs, implying diminished deformability of diabetic RBCs. The averaged values of mean membrane fluctuations for non-diabetic and diabetic RBCs are 55.1 ± 6.9 and 46.6 ± 4.5 nm, respectively. These observed diminutions of membrane fluctuation in diabetic cells are qualitatively in accordance with previous studies on decreased RBC deformability by diabetes based on micropipette aspiration^9^, laser diffraction ektacytometry^10, 42^ and techniques using optical tweezers^43^ and carbonate membrane filters^12^.

### Membrane fluctuations of RBCs in relation to HbA1c level

HbA1c reflects the degree of hyperglycaemia. HbA1c level is defined as a percentage of HbA1c, the majority of glycated Hbs in RBC, to the total Hb content in blood. Because HbA1c is formed over 120 days, non-enzymatic glycation process of Hb^44, 45^, the HbA1c level has been perceived as an effective means to estimate the mean glucose concentration of individuals for the previous three months^5, 46^.

Our results show a strong correlation between HbA1c and dynamic fluctuations in RBC membranes, as shown in Figure 4. The most noticeable alterations in diabetic RBCs can be found when the mean RBC membrane fluctuations of individuals are rearranged in the increasing order of the measured HbA1c levels. The mean membrane fluctuation of non-diabetic RBCs tends to decrease as the HbA1c level increases (Pearson correlation coefficient of −0.87 with a *p-*value of 0.02). In contrast, RBCs from diabetic patients exhibit significantly decreased membrane fluctuations, and the amplitude of dynamic membrane fluctuations are independent of the level of HbA1c. The mean values of membrane fluctuations for RBCs are 59.7 ± 6.2, 58.3 ± 4.9, 55.0 ± 5.4, 57.1 ± 7.4, 51.0 ± 6.8 and 49.6 ± 4.6 nm for healthy volunteers, and 45.1 ± 4.0, 46.9 ± 3.4, 48.8 ± 4.0, 47.6 ± 4.2, 43.0 ± 4.1 and 48.0 ± 5.0 nm for diabetic patients, in order of increasing HbA1c. This observed negative correlation between RBC membrane fluctuation and HbA1 is also conceptually in accordance with the report of a previous study, which reported a negative correlation between Hb1Ac levels and cell deformability indexes measured using a microchannel capillary model system^11^.

**Fig. 4.**
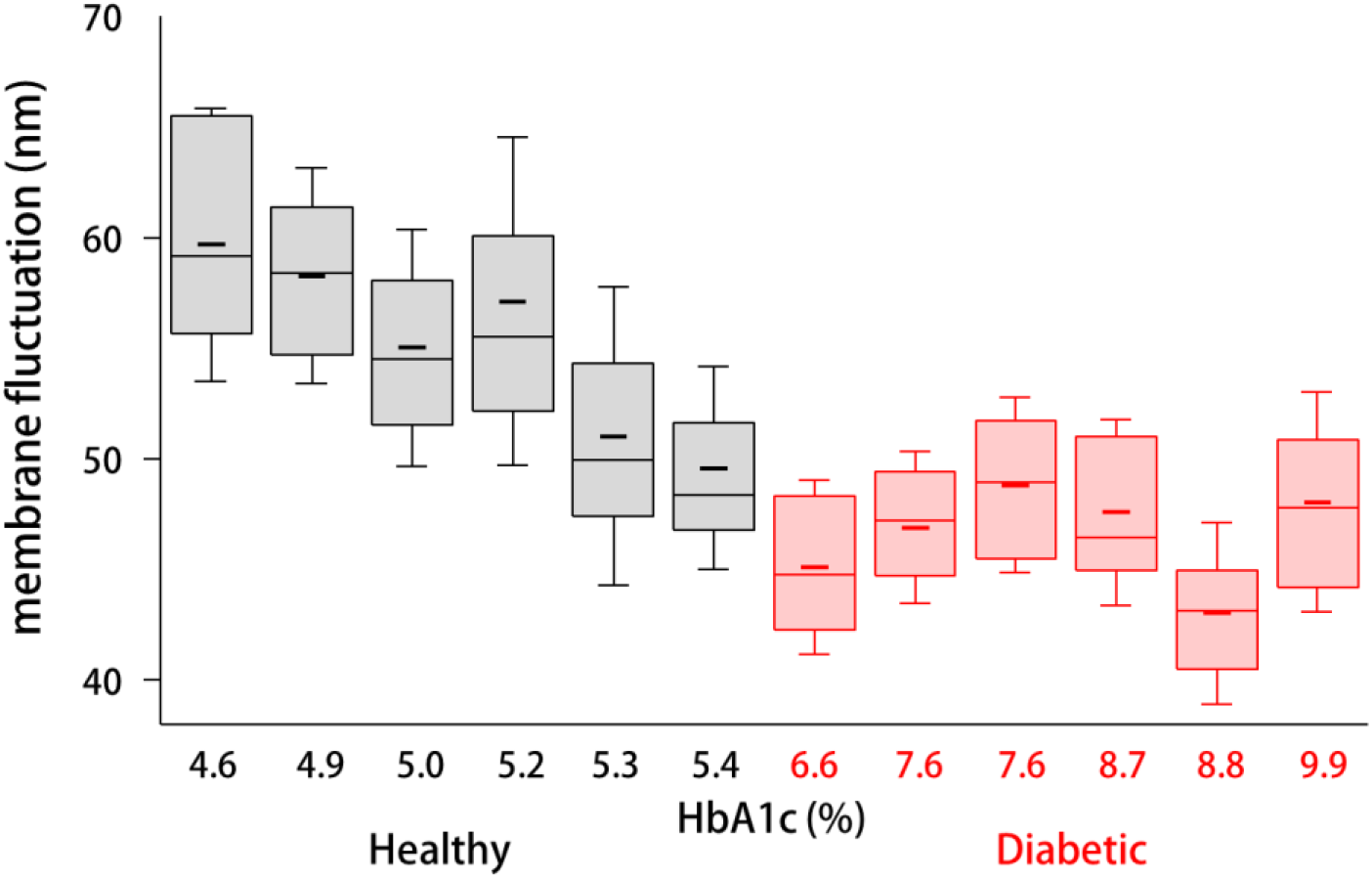
Box plots of membrane fluctuations for measured RBCs from six healthy controls and six diabetic patients in increasing order of HbA1c. Boxes show median values with upper and lower quartiles. Short horizontal lines and error bars in each box plot denote the mean values and standard deviations of individual cell measurements (n = approx. 40 per each group), respectively.

### Correlative analysis of individual RBCs

To fully exploit cellular alterations associated with diabetic complications, we performed correlative analyses among the retrieved parameters of individually measured non-diabetic and diabetic RBCs. As shown in Figure 5, RBC volume vs. Hb content, sphericity vs. membrane fluctuation and Hb concentration vs. membrane fluctuation are analysed and presented respectively.

**Fig. 5.**
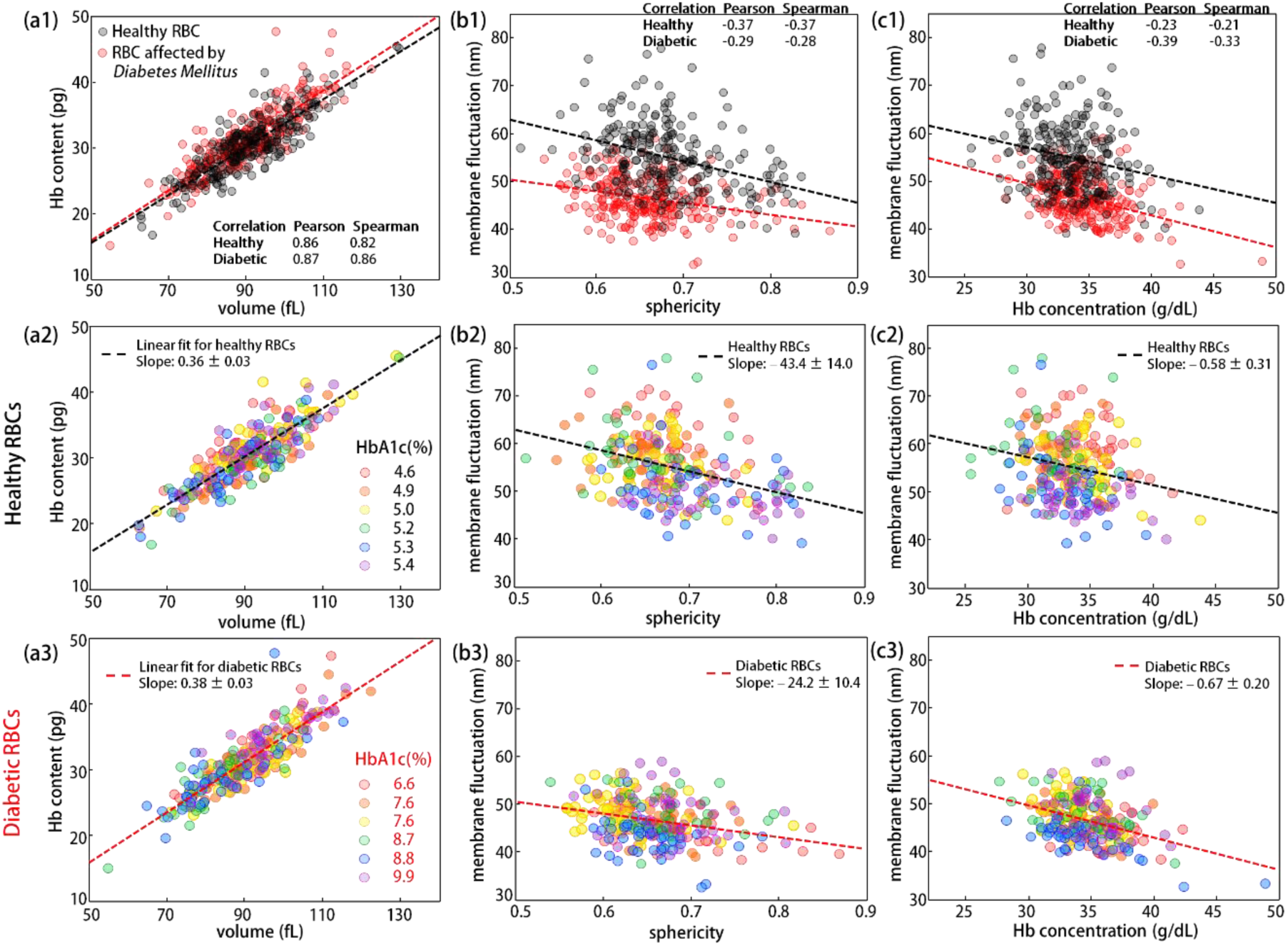
Scatter plots for various RBC parameters of non-diabetic and diabetic RBCs: (a1–a3) cell volume vs. Hb content, (b1–b3) sphericity vs. membrane fluctuation and (c1–c3) Hb concentration vs. membrane fluctuation. Each coloured circle denotes an individual RBC measurement. Scatter plots in the second (a2, b2 and c2) and third rows (a3, b3 and c3) represent RBC distributions of healthy and diabetic blood donors, respectively. RBCs from different individuals are presented in various colours according to HbA1c level. The dashed line in each figure represents a linear fit curve and the given slope covers 95% confidence interval. Pearson and Spearman coefficients for healthy and diabetic RBC groups are presented.

Statistical analysis using the linear regression model based on ANOVA shows that both healthy and diabetic RBCs share positive linear correlation between the cell volume and cytoplasmic Hb content, as shown in Figure 5(a1–a3). The slopes of the linearly fitted curves for non-diabetic and diabetic RBC populations are 36.4 ± 2.8 (with 95% confidence intervals) and 38.0 ± 2.8 g/dL, implying the respective intracellular Hb concentrations. Figures 5(a2) and (a3) present Hb content vs. volume correlation maps of non-diabetic and diabetic RBCs, respectively, and the circles denoting individual cell measurement are designated with different colours according to the HbA1c level of the blood donors. The observed larger slope for diabetic RBCs is in good agreement with our measurement of Hb concentration on individual RBCs.

Sphericity and membrane fluctuation of non-diabetic and diabetic RBCs have a negative correlation with both *p*-values less than 0.001, as shown in Figure 5(b1). Increased sphericity with reduced membrane fluctuations of RBCs associated with intracellular ATP depletion might account for the major portion of this overall negative correlation^32, 40^. The correlation maps show that RBCs from patients with diabetes exhibit significantly decreased mean membrane fluctuations compared to the healthy subjects in all ranges of sphericity. In Figures 5(b2) and (b3), the membrane fluctuation vs. sphericity correlation maps of non-diabetic and diabetic RBCs are respectively presented with the HbA1c-dependent coloured circle, so that cell distribution changes in relation to HbA1c level can be effectively addressed. In particular, it seems that the cell distribution of non-diabetic RBCs in Figure 5(b2) approaches to the lower-right end of the correlation map as HbA1c increases.

Also, the correlation map of Hb concentration vs. membrane fluctuation in Figure 5(c1) indicates that both non-diabetic and diabetic RBCs have a trend of exhibiting lower membrane fluctuations with higher Hb concentrations (*p-*values of less than 0.001 for both healthy and diabetic RBCs). Decreased deformability due to the elevated cytoplasmic viscosity of RBCs^37^ may account for this overall negative correlation between Hb concentration and membrane fluctuation, regardless of the RBC type.

## Discussion

### Morphological and biochemical alterations of diabetic RBCs

To our knowledge, our study is the first to present the simultaneous measurements of 3-D morphological, biochemical and mechanical properties of diabetic RBCs at the individual cell level. Previously, a few attempts have been made to investigate diabetic RBCs using quantitative phase imaging techniques. However, none of the previous work has comprehensively measured these parameters; all the previous works have been limited to 2-D phase imaging, where biochemical and morphological information are coupled to each other, and cannot be retrieved^47, 48^. In this work, the measured results show that diabetic RBCs exhibit statistically elevated cytoplasmic Hb concentration, Hb content and diminished sphericity when compared to those of non-diabetic RBCs. The non-enzymatic glycosylation, i.e. glycation, of Hb molecules^44, 45^ and membrane proteins^6^ by chronically elevated blood glucose content might account for the major portion of those alterations in diabetic RBCs. However, correlative analyses performed between those RBC parameters and the HbA1c level does not support clear dependency of morphological and biochemical parameters on HbA1c (Figure 5 and Supplementary Figure S1). Our experimental results also indicate that measured differences in mean values of Hb content and cell sphericity between non-diabetic and diabetic RBCs are comparable with variances of the corresponding parameters across the individual subjects. Accordingly, it seems that morphological and biochemical alteration in human RBCs by diabetes is not a radical and rapid process, but a mild and continual one.

### Diminished membrane fluctuation of diabetic RBCs: reduced deformability

Even between RBCs of similar values of sphericity or Hb concentration, considerable differences in membrane fluctuation of non-diabetic and diabetic RBCs were observed, as shown in Figures 5. (b1) and (c1). This implies that the mechanical alteration of the RBC membrane cortex is a leading cause of the diminished membrane fluctuations of diabetic RBCs. Widely known negative correlations between cell sphericity, Hb concentration and membrane fluctuation are also confirmed both in non-diabetic and diabetic RBCs. Mechanical alterations of diabetic RBC membranes are thought to be the consequences of the slow and irreversible glycation process of the cell membrane and intracellular proteins rather than the instant effects of high blood glucose. This is because previous research with the *in vitro* treatment of glucose on diluted blood does not significantly change the RBC deformability^13^. The observed negative correlation between the mean membrane fluctuation of RBCs and HbA1c level in Figure 4 also supports this view. Presumably, either an increase in membrane stiffness through glycation of band proteins comprising the spectrin network^10^ and membrane proteins^6^, or an accruing of peroxidation damages to the lipid membrane^7, 49^, or maybe both, account for the major portion of the reduced deformability of diabetic RBCs. Also, we found that no RBCs from diabetic patients exhibit mean membrane fluctuations greater than 50 nm. This observation suggests the possibility that the inherent fluctuating motion of RBCs can be a viable option for assisting the diagnosis or prevention of diabetic mellitus because the exploitation of exogenous agents or complicated techniques are not required.

## Conclusion

We have quantitatively and non-invasively characterised the morphological, biochemical and mechanical alterations of individual RBCs by diabetes mellitus. The HbA1c levels of subjects, which reflect one’s mean blood glucose concentration of the previous three months, were also determined using HbA1c analysers. Accordingly, correlative analyses among the retrieved individual RBC parameters and HbA1c level were performed to search for unique features which cannot be addressed by conventional techniques incapable of simultaneous measurement of those parameters.

Our optical measurement reports slightly elevated cytoplasmic Hb concentration and content, and reduced sphericity of diabetic RBCs compared to the control RBCs. Taking into account the fact that observed differences in RBC parameters between healthy blood donors and diabetic patients are not much larger than variances of those parameters among individual people, it is difficult to see that morphological and biochemical properties of human RBCs are altered by diabetes significantly enough to be used for the purpose of diagnosis. The most dramatic alteration in diabetic RBCs can be found in their membrane fluctuations; diabetic RBCs exhibit membrane fluctuations significantly lower than those of non-diabetic RBCs. Successively performed correlative analyses further show that healthy donors tend to have RBCs of higher sphericity and lower membrane fluctuation, with a higher HbA1c value. These observations imply that hyperglycaemia associated with diabetes severely impairs RBC deformability by remodelling mechanical properties of the cell membrane.

With capabilities enabling quantitative measurements of individual microscopic cells under biocompatible conditions, we envision that QPI methodologies could shed light on unresolved areas of diabetes mellitus. In particular, a recently developed miniaturised QPI unit^50-52^ can convert conventional bright field microscopy into a QPI system by simply attaching the unit to a microscope body with light source adjustment. Furthermore, QPI techniques employing DMD-based illumination schemes^53^ or graphic-processing units^54^ allow real-time visualisation of a 3-D microscopic object, and are expected to become an attractive imaging tool for investigating live cell dynamics.

## Materials and Methods

### Blood sample preparation and ethics statement

3 mL of blood was collected from six patients with type 2 diabetes mellitus by venipuncture and transferred into EDTA anticoagulant tubes at Asan Medical Centre, Seoul, Republic of Korea. CBC tests were instantly performed using an XE-2100 automated haematology analyser (Sysmex Co., Kobe, Japan), from which mean corpuscular Hb, mean corpuscular Hb concentration and mean corpuscular volume with RBC distribution width of individual diabetic patients were obtained. Independently, HbA1c was determined based on high-performance liquid chromatography using a TOSOH G8 analyser (Tosoh Co., Tokyo, Japan). For optical measurement, blood samples were delivered to a certified laboratory (IRB project: KH2015-37) at KAIST (Korea Advanced Institute of Science and Technology), Daejeon, Republic of Korea, within four hours of blood collection. All blood gathering protocols performed at Asan medical centre were approved by the ethics committee of the University of Ulsan, Ulsan, Republic of Korea (IRB project: #IRB-13-90). For the purpose of comparison, control blood was drawn by venipuncture from six healthy blood donors with the same blood collection protocols at a health clinic (the Pappalardo Centre at KAIST. A XT-2000i (Sysmex Co., Kobe, Japan) haematology analyser and a Cobra Integra 800 chemistry analyser (Roche Diagnostics, Risch-Rotkreuz, Switzerland) were used for the CBC test and determination of HbA1c, respectively. All the experimental protocols for the control blood donors were approved by the institutional review board of KAIST (IRB project: KH2015-37). Blood samples were stored in a refrigerator of 4°C and optically measured using cDOT within 24 hours of blood gathering. In relation to the storage process, a recent study reported a result of invariances in mean membrane fluctuation of stored RBCs at 4°C within 2 or 3 days after blood collection^55^. All the blood samples were collected as part of a regular course of patient care in Asan Medical Centre, and we selected patients who had given informed consent for using their archival tissues for genetic testing. All data was de-identified.

### Optical measurement on individual RBCs

To optically measure the individual RBCs, 3 μL of collected blood was diluted by a factor of 300 with Dulbecco’s phosphate buffered saline (Gibco®, New York, U.S.A.). Diluted blood suspensions were then sandwiched between two coverslips (24 × 50 mm, C024501, Matsunami, Ltd, Japan) and loaded on a sample stage of the inverted microscope body. After 20 min of waiting for cell settlements on a bottom coverslip, 40 RBCs for each healthy donor or diabetic patient were measured. In measuring RBCs, only gently sedimented RBCs of discocytes were selected. After excluding erroneously measured RBCs, 40, 40, 40, 38, 39 and 40 control RBCs of six healthy donors, and 40, 40, 40, 40, 40 and 40 diabetic RBCs of six diabetic patients were analysed respectively for the study of diabetic effects on characteristics of individual human RBCs.

### Statistical analysis

Specified *p*-values in the manuscript were calculated based on a two-tailed student’s t-test comparing mean RBC parameters between control and diabetes RBC groups. Retrieved RBC parameters were given in the manuscript as mean ± SD. In the correlative analyses, an analysis of variance (ANOVA) based linear regression model was employed to obtain linear fit curves for both control and diabetic RBCs. All the numbers following a ± sign in the linear regression model cover a 95% confidence interval. To further investigate the correlative nature of various RBC parameters along the HbA1c level, Pearson (linear correlation) and Spearman (monotonous correlation) correlation coefficients were calculated. Throughout the manuscript, a *p*-value of less than 0.05 was regarded as statistically significant.

### Common-path Diffraction Optical Tomography (cDOT)

For simultaneous measurements of both 3-D RI tomograms and 2-D membrane fluctuations of RBCs, we employed cDOT^24^. A diode-pumped solid-state laser (λ = 532 nm, 50 mW, Cobolt Co., Solna, Sweden) was used as a light source. The laser impinges on a sample loaded on the stage of an inverted microscope with various illumination angles, which is controlled using a dual-axis galvanometer mirror (GM1; GVS012/M, Thorlabs, U.S.A.). A condenser objective lens [UPLFLN 60×, numerical aperture (NA) = 0.9, Olympus Inc., San Diego, CA, U.S.A.] was used for illuminating a sample with a broad range of impinging angles, and the high-NA objective lens (PLAPON 60×, oil immersion, N.A. = 1.42, Olympus Inc., San Diego, CA, U.S.A.) was used to collect diffracted light from a sample. The other galvanometer mirror (GM2), located at the conjugated plane of the sample, was synchronised with GM1 so that the laser beam after GM2 always followed the same path regardless of the laser illumination angle.

The rest of cDOT constitutes common-path laser interferometry. Using a diffraction grating (92 grooves/mm, #46-072, Edmund Optics Inc., NJ, U.S.A.), the laser beam after GM2 was divided into various diffracted orders. The brightest 0^th^ order diffracted beam was then spatially filtered by passing through a small pinhole (Ø25 μm, P25S, Thorlabs, U.S.A.) located at the Fourier plane, and served as a reference arm in off-axis interferometry. The 1^st^ order diffracted beam containing the sample information served as a sample arm. Then, the spatially modulated interferograms of the sample were formed at the image plane, which was recorded by the high-speed sCMOS camera (Neo sCMOS, ANDOR Inc., Northern Ireland, U.K.). The total magnification of the system was 250×, contributed from both the 60× imaging objective lens and the additional telescopic 4-*f* imaging system. The field of view of cDOT was 14.3 × 13.31 μm^2^, which corresponded to 528 × 512 camera pixels with a pixel size of 6.5 μm

From the captured interferograms, the complex optical fields, consisting of amplitude and phase information, were retrieved by using the field-retrieval algorithm^29^. Then, from a set of the retrieved complex optical fields obtained at 300 different illumination angles, a 3-D RI tomogram of the sample *n*(*x*, *y*, *z*) was reconstructed employing optical diffraction tomography^26^. To fill the missing cone information caused by the limited NAs of the lenses, the regularisation algorithm based on non-negativity criteria was employed^56^. Detailed information about optical diffraction tomography including the Matlab code can be found elsewhere^27^. For the visualisation of the measured 3-D RI tomograms, commercial software (Tomostudio, Tomocube, Inc., Daejeon, Republic of Korea) was used.

To measure 2-D height profiles and membrane fluctuations of RBCs, the laser illumination was set to be perpendicular on the sample. Consecutive interferograms were then continuously recorded with the high-speed camera at a frame rate of 125 Hz for 2.4 sec, from which 2-D phase delay maps Δ*ϕ* (*x*, *y*, *t*) were retrieved using the field retrieval algorithm. 2-D RBC height profiles *h*(*x*, *y*, *t*) were then calculated according to the following relation: *h*(*x*, *y*, *t*) = [λ/(2*π*·〈Δ*n*〉)]·Δ*ϕ* (*x*, *y*, *t*).

### Analysis procedures for retrieving RBC parameters

All the retrieved RBC parameters include morphological (volume *V*, surface area *S* and sphericity *SI*), biochemical (Hb concentration [Hb] and Hb content) and mechanical (membrane fluctuation σ_h_) properties of individual RBCs.

Of the morphological parameters, *V* and *S* of an individual RBC can be directly obtained from the reconstructed 3-D RI tomogram. Sphericity, *SI*, a measure of sphere resemblance, is then calculated as a normalised volume-to-surface area ratio as follows: *SI* = (6*V*)^2/3^π^1/3^/*S*.

To obtain [Hb] from tomographic measurements of RBCs, it needs to be noted that the RI difference between the cytoplasm and the surrounding medium is linearly proportional to the concentrations of intracellular non-aqueous solutes with a proportionality coefficient, α, which is a so-called refraction increment^30^. This relation can be written as follows: 〈Δ*n*〉 = α[Hb], where 〈Δ*n*〉 = 〈*n*(*x*, *y*, *z*)〉_spatial_ − *n*_*m*_, *n*(*x*, *y*, *z*) is a spatial RI distribution of a sample, and *n*_*m*_ is a medium RI. For mixtures of various chemicals, α is determined as a linear combination of α values assigned for each chemical with weight coefficients of respective concentrations. The practical issue is whether there are detectable differences in values of α between RBCs from healthy controls and ones from diabetic patients. It has been known that the most prominent alteration in cell components by hyperglycaemia is the glycosylation of intracellular proteins. In particular, when it comes to RBCs, Hb constitutes more than 90% of total RBC proteins, and the percentage of HbA1c to total Hb content can be elevated at most by 6 ~ 7% in diabetic patients, as compared with normal HbA1c range^5^. Accordingly, expected changes in value of α by diabetes are thought to be less than 0.001 mL/g because most proteins exhibit similar values of α - at most 5% difference in α with each other^57, 58^. Hence, α was set to have 0.18 mL/g^59^ for RBCs from both healthy controls and ones from diabetic patients. Then, cytoplasmic Hb contents can be obtained by multiplying *V* by [Hb].

In this paper, the 2-D membrane fluctuation map, σ_h_ (*x*, *y*), is defined as a temporal standard deviation of mean height profiles of RBCs, *h*(*x*, *y*, *t*), and can be written as:

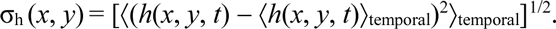

The membrane fluctuation, σ_h_, is then a spatial average of the 2-D fluctuation map, σ_h_ = 〈σ_h_ (*x*, *y*)〉_spatial_ over projection cell area, and serves as the intuitive measure of RBC deformability.

## Acknowledgements

This work was supported by KAIST, and the National Research Foundation of Korea (2015R1A3A2066550, 2014K1A3A1A09063027, 2012-M3C1A1-048860, 2014M3C1A3052537) and Innopolis foundation (A2015DD126).

## Author contributions

SL, SJ, YP conceived and designed the experimental idea. SL, HP performed the experiments and analyzed the data. SL, HP, KK, SJ, and YP wrote the manuscript.

## References

1 Forbes, J. M. & Cooper, M. E. Mechanisms of diabetic complications. Physiological reviews 93, 137–188 (2013).

2 Turchetti, V. et al. Evaluation of erythrocyte morphology as deformability index in patients suffering from vascular diseases, with or without diabetes mellitus: correlation with blood viscosity and intra‐erythrocytic calcium. Clinical hemorheology and microcirculation 18, 141–149 (1998).

3 Davidson, R., Evan-Wong, L. & Stowers, J. The mean red cell volume in diabetes mellitus. Diabetologia 20, 583–584 (1981).

4 Piagnerelli, M. et al. Assessment of erythrocyte shape by flow cytometry techniques. Journal of clinical pathology 60, 549–554 (2007).

5 Rohlfing, C. L. et al. Defining the relationship between plasma glucose and HbA1c analysis of glucose profiles and HbA1c in the Diabetes Control and Complications Trial. Diabetes care 25, 275–278 (2002).

6 Miller, J. A., Gravallese, E. & Bunn, H. F. Nonenzymatic glycosylation of erythrocyte membrane proteins. Relevance to diabetes. Journal of Clinical Investigation 65, 896 (1980).

7 Jain, S. K. Hyperglycemia can cause membrane lipid peroxidation and osmotic fragility in human red blood cells. Journal of Biological Chemistry 264, 21340–21345 (1989).

8 Jain, S. K., Levine, S. N., Duett, J. & Hollier, B. Elevated lipid peroxidation levels in red blood cells of streptozotocin-treated diabetic rats. Metabolism 39, 971–975 (1990).

9 McMillan, D. E., Utterback, N. G. & La Puma, J. Reduced erythrocyte deformability in diabetes. Diabetes 27, 895–901 (1978).

10 Schwartz, R. S., Madsen, J. W., Rybicki, A. C. & Nagel, R. L. Oxidation of spectrin and deformability defects in diabetic erythrocytes. Diabetes 40, 701–708 (1991).

11 Tsukada, K., Sekizuka, E., Oshio, C. & Minamitani, H. Direct measurement of erythrocyte deformability in diabetes mellitus with a transparent microchannel capillary model and high-speed video camera system. Microvascular research 61, 231–239 (2001).

12 Ernst, E. & Matrai, A. Altered red and white blood cell rheology in type II diabetes. Diabetes 35, 1412–1415 (1986).

13 Resmi, H., Akhunlar, H., Temiz Artmann, A. & Güner, G. In vitro effects of high glucose concentrations on membrane protein oxidation, G‐actin and deformability of human erythrocytes. Cell biochemistry and function 23, 163–168 (2005).

14 Popescu, G. Quantitative Phase Imaging of Cells and Tissues. (McGraw-Hill Professional, 2011).

15 Lee, K. et al. Quantitative Phase Imaging Techniques for the Study of Cell Pathophysiology: From Principles to Applications. Sensors 13, 4170–4191 (2013).

16 Park, Y. et al. Refractive index maps and membrane dynamics of human red blood cells parasitized by Plasmodium falciparum. Proceedings of the National Academy of Sciences 105, 13730–13735 (2008).

17 Diez-Silva, M. et al. Pf155/RESA protein influences the dynamic microcirculatory behavior of ring-stage Plasmodium falciparum infected red blood cells. Scientific reports 2 (2012).

18 Chandramohanadas, R. et al. Biophysics of malarial parasite exit from infected erythrocytes. PLoS One 6, e20869, doi:10.1371/journal.pone.0020869 (2011).

19 Kim, Y. et al. Profiling individual human red blood cells using common-path diffraction optical tomography. Scientific reports 4 (2014).

20 Yoon, J. et al. Label-free characterization of white blood cells by measuring 3D refractive index maps. Biomed Opt Express 6, 3865–3875 (2015).

21 Jourdain, P. et al. Determination of transmembrane water fluxes in neurons elicited by glutamate ionotropic receptors and by the cotransporters KCC2 and NKCC1: a digital holographic microscopy study. The Journal of Neuroscience 31, 11846–11854 (2011).

22 Byun, H. et al. Optical measurement of biomechanical properties of individual erythrocytes from a sickle cell patient. Acta Biomaterialia (2012).

23 Jung, J. et al. Optical characterization of red blood cells from individuals with sickle cell trait and disease in Tanzania using quantitative phase imaging. arXiv preprint arXiv:1604.06796 (2016).

24 Kim, Y. et al. Common-path diffraction optical tomography for investigation of three-dimensional structures and dynamics of biological cells. Optics express 22, 10398–10407 (2014).

25 Shin, S. et al. in SPIE BiOS. 933629-933629-933626 (International Society for Optics and Photonics).

26 Wolf, E. Three-dimensional structure determination of semi-transparent objects from holographic data. Optics Communications 1, 153–156 (1969).

27 Kim, K. et al. High-resolution three-dimensional imaging of red blood cells parasitized by *Plasmodium falciparum* and *in situ* hemozoin crystals using optical diffraction tomography. J. Biomed. Opt. 19, 011005–011012 (2014).

28 Kim, K. et al. Optical diffraction tomography techniques for the study of cell pathophysiology. arXiv preprint arXiv:1603.00592 (2016).

29 Debnath, S. K. & Park, Y. Real-time quantitative phase imaging with a spatial phase-shifting algorithm. Optics letters 36, 4677–4679 (2011).

30 Barer, R. Determination of dry mass, thickness, solid and water concentration in living cells. Nature 172, 1097–1098 (1953).

31 Popescu, G. et al. Optical imaging of cell mass and growth dynamics. American Journal of Physiology - Cell physiology 295, C538–544, doi:10.1152/ajpcell.00121.2008 (2008).

32 Park, Y. et al. Metabolic remodeling of the human red blood cell membrane. Proceedings of the National Academy of Sciences 107, 1289 (2010).

33 Shaked, N. T., Satterwhite, L. L., Telen, M. J., Truskey, G. A. & Wax, A. Quantitative microscopy and nanoscopy of sickle red blood cells performed by wide field digital interferometry. Journal of biomedical optics 16, 030506 (2011).

34 Lee, S. Y., Park, H. J., Best-Popescu, C., Jang, S. & Park, Y. K. The Effects of Ethanol on the Morphological and Biochemical Properties of Individual Human Red Blood Cells. PloS one 10, e0145327 (2015).

35 Park, H. et al. Characterizations of individual mouse red blood cells parasitized by Babesia microti using 3-D holographic microscopy. Scientific Reports 5, doi:10.1038/Srep10827 (2015).

36 Cranston, H. A. et al. Plasmodium falciparum maturation abolishes physiologic red cell deformability. Science 223, 400–403 (1984).

37 Mohandas, N. & Chasis, J. in Seminars in hematology. 171–192.

38 Kim, Y., Kim, K. & Park, Y. in Blood Cell - An Overview of Studies in Hematology (ed Terry E. Moschandreou) Ch. 10, 167–194 (INTECH, 2012).

39 Hosseini, P. et al. Cellular normoxic biophysical markers of hydroxyurea treatment in sickle cell disease. Proceedings of the National Academy of Sciences, 201610435 (2016).

40 Park, H. et al. Alterations in cell surface area and deformability of individual human red blood cells in stored blood. arXiv preprint arXiv:1506.05259 (2015).

41 Park, H. et al. Three-dimensional refractive index tomograms and deformability of individual human red blood cells from cord blood of newborn infants and maternal blood. J Biomed Opt 20, 111208, doi:10.1117/1.jbo.20.11.111208 (2015).

42 Shin, S., Ku, Y., Babu, N. & Singh, M. Erythrocyte deformability and its variation in diabetes mellitus. Indian journal of experimental biology 45, 121 (2007).

43 Smart, T. J. et al. in SPIE Nanoscience+ Engineering. 954825-954825-954827 (International Society for Optics and Photonics).

44 Bunn, H. F., Haney, D. N., Gabbay, K. H. & Gallop, P. M. Further identification of the nature and linkage of the carbohydrate in hemoglobin A 1c. Biochemical and biophysical research communications 67, 103–109 (1975).

45 Bunn, H. F., Gabbay, K. H. & Gallop, P. M. The glycosylation of hemoglobin: relevance to diabetes mellitus. Science 200, 21–27 (1978).

46 Gallagher, E. J., Le Roith, D. & Bloomgarden, Z. Review of hemoglobin A1c in the management of diabetes. Journal of diabetes 1, 9–17 (2009).

47 Talaykova, N., Kalyanov, A., Lychagov, V., Ryabukho, V. & Malinova, L. in 1st International Conference. 90320F-90320F-90325 (International Society for Optics and Photonics).

48 Doblas, A. et al. Diabetes screening by telecentric digital holographic microscopy. Journal of microscopy (2015).

49 Jain, S. K., McVie, R., Duett, J. & Herbst, J. J. Erythrocyte membrane lipid peroxidation and glycosylated hemoglobin in diabetes. Diabetes 38, 1539–1543 (1989).

50 Lee, K. & Park, Y. Quantitative phase imaging unit. Optics letters 39, 3630–3633 (2014).

51 Kim, K. et al. Diffraction optical tomography using a quantitative phase imaging unit. Optics letters 39, 6935–6938 (2014).

52 Baek, Y., Lee, K., Yoon, J., Kim, K. & Park, Y. White-light quantitative phase imaging unit. Optics express 24, 9308–9315 (2016).

53 Shin, S., Kim, K., Yoon, J. & Park, Y. Active illumination using a digital micromirror device for quantitative phase imaging. Optics letters 40, 5407–5410 (2015).

54 Kim, K., Kim, K. S., Park, H., Ye, J. C. & Park, Y. Real-time visualization of 3-D dynamic microscopic objects using optical diffraction tomography. Optics Express 21, 32269–32278 (2013).

55 Park, H. et al. Measuring cell surface area and deformability of individual human red blood cells over blood storage using quantitative phase imaging. Scientific Reports 6 (2016).

56 Lim, J. et al. Comparative study of iterative reconstruction algorithms for missing cone problems in optical diffraction tomography. Optics Express 23, 16933–16948, doi:10.1364/Oe.23.016933 (2015).

57 Barer, R. & Joseph, S. Refractometry of living cells part I. Basic principles. Quarterly Journal of Microscopical Science 3, 399–423 (1954).

58 Stramski, D. Refractive index of planktonic cells as a measure of cellular carbon and chlorophyll a content. Deep Sea Research Part I: Oceanographic Research Papers 46, 335–351 (1999).

59 Park, Y., Yamauchi, T., Choi, W., Dasari, R. & Feld, M. S. Spectroscopic phase microscopy for quantifying hemoglobin concentrations in intact red blood cells. Optics letters 34, 3668–3670 (2009).

